# Best practice mass photometry: A guide to optimal single molecule mass measurement

**DOI:** 10.1101/2024.12.03.624087

**Authors:** Jiří Kratochvíl, Raman van Wee, Dan Loewenthal, Jan Christoph Thiele, Jack Bardzil, Kishwar Iqbal, Stephen Thorpe, Philipp Kukura

## Abstract

Mass photometry (MP) has emerged as a powerful approach to study biomolecular structure, dynamics and interactions. The capabilities of the method ultimately hinge on the ability to accurately measure the tiny optical contrast generated by individual molecules landing at a glass-water interface, which enables mass-resolved quantification of biomolecular mixtures. Ideally, this capability is only limited by shot noise inherent to photon detection, but in practice depends on additional parameters and details of the assay. Here, we focus on the key parameters affecting MP performance, and present simple steps that can be taken to achieve optimal MP performance in terms of mass resolution, quantitative detection limit and analyte concentration range without compromising the ease and simplicity of the technique.

## Introduction

Mass photometry (MP) is a label-free method for the solution-based mass measurement of single biomolecules^1^. Due to its simplicity, speed and unique information content, MP has been adopted widely to study a large variety of phenomena, such as protein oligomerisation, adeno-associated virus assembly, protein-DNA and antibody-antigen interactions, among many others^2,3,4,5^. In addition to the original implementation relying on non-specific adsorption of the analyte to a bare glass surface, dynamic measurements of tracking the analyte movement on the surface of lipid-bilayers are also possible^6,7^. Previous protocols for MP have focused on the basics of performing an MP measurement with a focus on binding affinities^8,9^.

Standard MP experiments require as little as the addition of a few µl of sample to a microscope coverglass, similar to a nanodrop measurement. The resulting data provide immediate information on sample composition, resolved by molecular mass. Owing to its simplicity, MP is extremely robust and broadly applicable. At the same time, minimal changes to the assay can dramatically improve the final data quality. By systematically exploring the influence of noise sources affecting single molecule measurement precision, we illustrate and define a workflow that mitigates potential pitfalls and thereby enables easy optimization of MP measurements. As a result, we enable the user to routinely obtain the best possible data quality without affecting the simplicity and speed of MP measurements.

### Principle of MP

The basic detection principle of MP rests on quantifying the difference in the reflectivity of an interface in the presence and absence of a biomolecule. Most commonly, that change originates from the interference between light scattered by the biomolecule binding to a glass-water interface, and light reflected by the same interface (**Fig. 1a**). The use of an amplitude mask selectively attenuates reflected light without affecting the scattered light, enhancing the optical contrast and enabling much higher power density at the sample without saturating the detector, thereby improving the experimentally achievable signal-to-noise ratio (SNR) for a given exposure time^10^.

**Figure 1.**
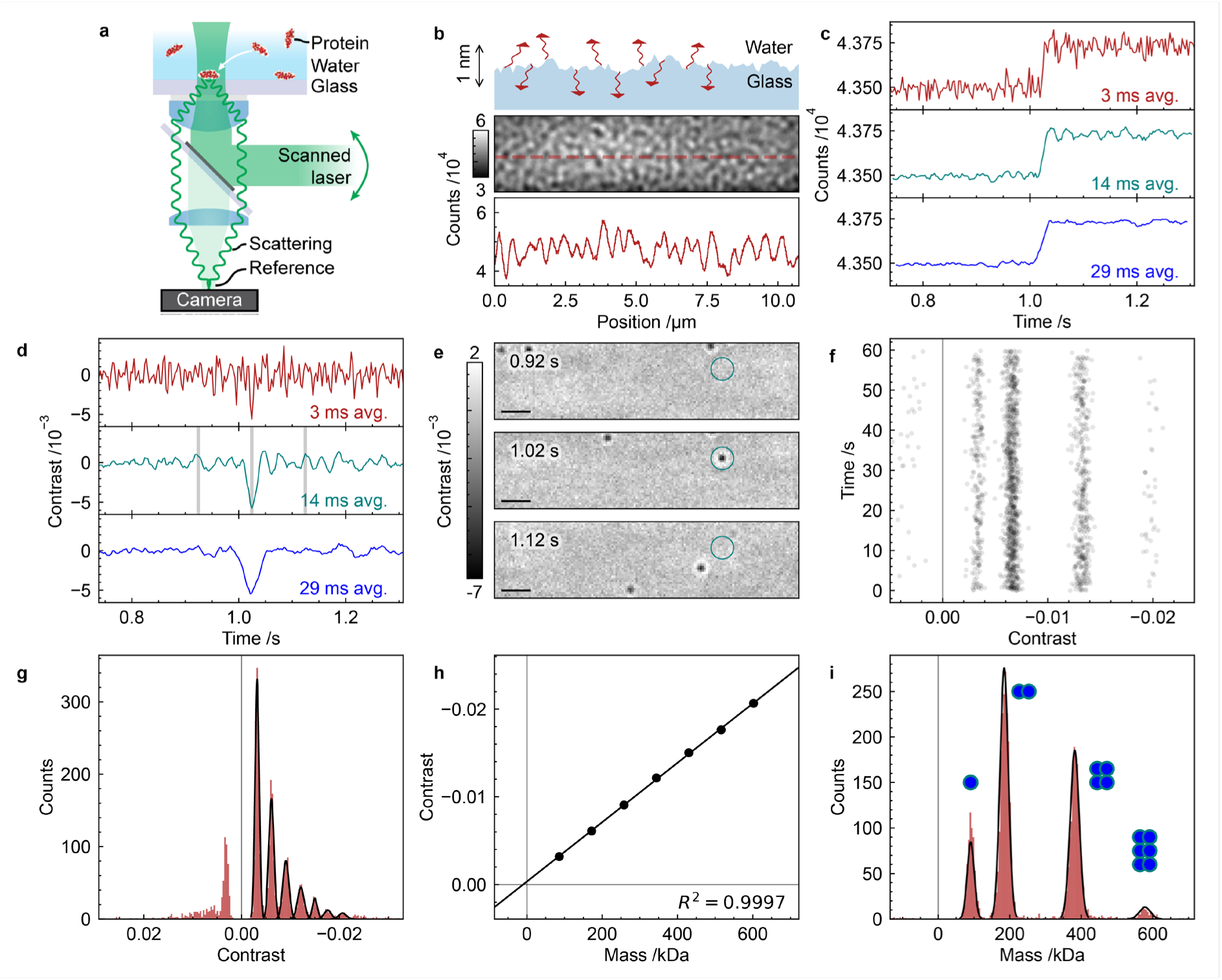
Principles of Mass Photometry. (**a**) Measurement principle: Single proteins or complexes bind to the surface and become immobile. The light scattered by the protein is quantified to infer its mass. (**b**) The raw data is dominated by the static scattering from the glass roughness at the glass water interface. (**c**) Measurement of a diluted protein sample, Dyn1 ΔPRD. Intensity time trace of the landing event for a 180 kDa species. The lower two panels are smoothed with moving average. (**d**) Corresponding traces in the ratiometric data calculated with different window sizes. (**e**) Ratiometric frames at different time points (window size 14 ms, Scale bar 1 μm). (**f**) Scatter plot of the extracted contrast and the landing time. (**g**) Contrast distribution for the calibration protein MFP1 (**h**) The conversion between mass and contrast is determined based on the fits to the measured contrast and known masses of the calibration sample MFP1. (**i**) The mass distribution of the protein Dyn1 ΔPRD shows its oligomeric nature.

The raw image of the interface captured by the camera is dominated by scattering caused by the nanoscopic roughness of the glass substrate, which results in a speckle-like pattern (**Fig. 1b**)^11,12^. Depending on the illumination wavelength and mask strength used, the resulting intensity variation is on the order of 10-30% of the reflected light intensity peak-to-peak. Binding of a 180 kDa protein to this surface results in a small change in the reflectivity, in this case on the order of 0.5%, the visibility of which improves with temporal averaging by reducing shot noise induced fluctuations in the background light intensity (**Fig. 1c**). While this step change is easily visible when zoomed in on a single camera pixel, it is undetectable in the raw image, because the signal magnitude (0.5%) is 1-2 orders of magnitude smaller than the image speckle caused by the glass surface roughness.

The lack of visibility can be addressed by averaging out the background image fluctuations by computing ratiometric images as a function of time, which consists of the ratio between two windows of multiple frames averaged to reduce shot noise-induced background fluctuations (**Fig. 1c**). This approach converts a small step change on top of a large background into one on top of shot noise-limited background, enhancing visibility and simplifying detection (**Fig. 1d**). The normalized contrast is given by Eq. 1

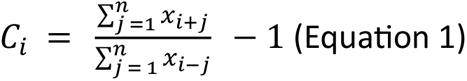

where *n* is the window size in frames and *x* the raw camera frames recorded by the camera. Computing the ratiometric image for a binding event at the maximum of this trace yields an image of the binding event, whose visibility is strongly affected by the degree of spatial and temporal binning, which in turn affects shot noise-induced background noise (**Fig. 1e**).

In an experiment where the analyte is present in solution, proteins continuously bind to the glass interface. Given sufficient single molecule contrast measurement precision, the contrast variation from event to event is small enough, such that species of different mass can be resolved, resulting in clear stripes when plotting the detected contrast as a function time (**Fig. 1f**). Integrating these events into a contrast histogram yields baseline-resolved peaks at integer multiples of the monomer contrast (**Fig. 1g**). To convert from contrast to mass a known calibrant protein is measured initially. Plotting the center of mass of each peak vs the known molecular mass of the different calibrant oligomers yields an accurate mass-to-contrast relationship (**Fig. 1h**). This calibration can then be applied to unknown samples, which together with the high achievable mass resolution, enables identification and quantification of biomolecular mixtures by mass (**Fig. 1i**).

The key performance metrics for MP measurements are: (1) Mass accuracy; the difference between the measured and expected mass; (2) Mass resolution; the smallest mass difference for which two species in a mixture can be resolved; (3) Quantification; to which degree the measured molecular counts are proportional to the abundance of each species in the mixture. For optimal measurements, these values have reached 2% accuracy, 20 kDa resolution and a detection limit on the order of 40 kDa.^1^ Here, we identify and quantify different noise sources, show how they affect these key metrics and thereby provide guidelines for how to generally and easily achieve optimal data quality.

### Repeatability and dynamic range

To characterise the reproducibility of MP measurements, we consecutively measured eight replicates of a mass calibration protein standard (**Fig. 2a**). To achieve maximum reproducibility in the total counts, it is important to ensure that the incubation time in the sample tube and the time between sample addition and the beginning of the measurement is consistent. Under such conditions, the total event count across eight replicates fluctuates by about ±15% around the mean of the total event counts. This number can differ depending on how and where the analyte is added and convection within the droplet. The monomer:dimer ratio, exhibits a smaller fluctuation on the order of ± 5% (**Fig. 2b**). This ratio is less sensitive to how the sample is added and rather depends on sample concentration and incubation time through the relevant interaction affinities.

**Figure 2.**
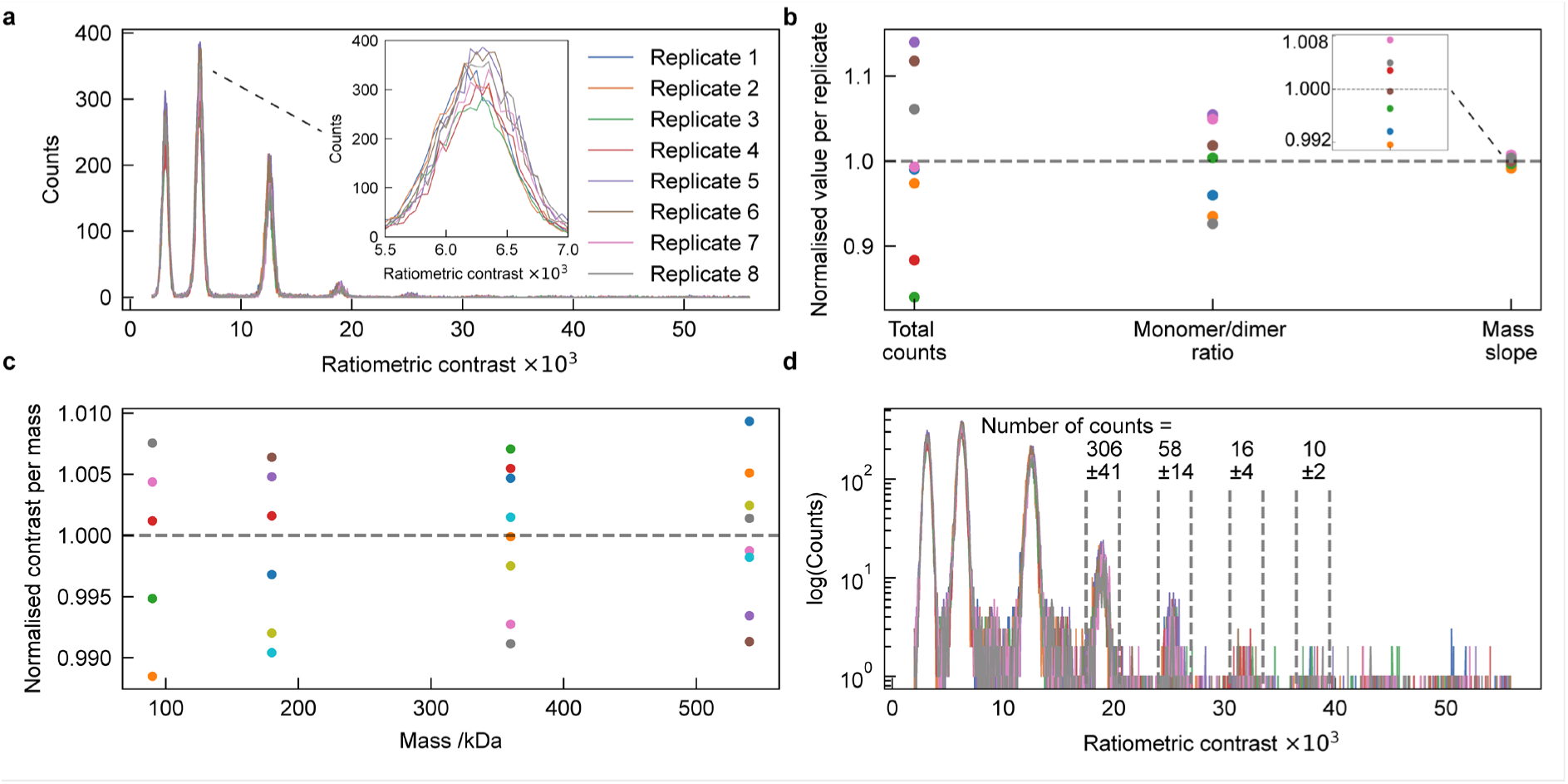
Measurement reproducibility. Dataset of 8 consecutively measured replicates of Dyn1 ΔPRD. (**a**) Detected events vs ratiometric contrast histograms, with a magnification of a single ratiometric contrast peak (inset). (**b**) Reproducibility metrics; consecutive measurements conducted according to our protocol can yield a total events count spread of about 30%, an oligomeric ratio spread of about 10% and a fitted mass slope spread of about 2%. (**c**) Normalized fitted ratiometric contrast values for the different oligomeric states; the fitted peak ratiometric contrast values for all oligomeric states vary by about 2% between replicates. (**d**) Log-scaled detected events vs ratiometric contrast histogram that can highlight peaks that are hard to see on linear-scale plot.

The retrieved mass-to-contrast slope from a fit as shown in **Fig. 1h** can vary by as a little as ±1% when care is taken in terms of sample addition (**Fig. 2b, see procedure**). This observation is confirmed when examining the ratiometric contrast of the different oligomers (**Fig. 2c**), which is again limited to 2% of the oligomer mass. This variation limits the smallest changes detectable by MP, where complete analyte population shifts from mass m to mass m+2%, which can be as low as 1 kDa^1^. We emphasise the difference here between measurement precision, which is sufficient for visualising small molecule binding, and resolution, which requires the simultaneous and independent observation of two species. In general, the repeatability in terms of mass measurement is most sensitive to the sample focus, which can be reliably optimized by maximising the optical contrast of the glass roughness in the raw image.

An important aspect of MP that has been largely neglected to date is the dynamic range in terms of relative abundance of species present in the mixture. Given that single biomolecules or their complexes are detected in a digital fashion, the dynamic range is in principle only limited by measurement time. For the example shown here, a linear representation of the data reveals monomer, dimer, tetramer, hexamer and small amounts of octamer. Plotting the same data on a log scale (**Fig. 2d**) shows clear signatures of decamers and dodecamers, as well as low counts of even larger species. As previously, the reproducibility of these oligomeric ratios is very high. In practice, the dynamic range is often limited by impurities in the sample, but if these can be avoided and total counts exceed 10^4^, it can reach >>10^3^, as shown by the reproducible detection of a species with only 10 counts in a background of 10^4^ counts (**Fig. 2d**).

### The impact of detection threshold, averaging time and field of view

In the absence of other noise sources, the ability to detect and quantify individual biomolecules by MP is only limited by shot noise-induced fluctuations of the imaging background from reflected light at the glass-water interface. For shot noise, the expected fluctuation for the detection of N photoelectrons per pixel is √N, thus the magnitude of background fluctuations is given by √N/N = 1/√N. Detecting more photoelectrons per unit time thus reduces the image noise and improves detection and quantification of individual biomolecular events.

The incident power is limited first by the damage threshold of the optical components and sample, and second by the finite full well depth and frame rate of the detector. Increasing the temporal averaging window means more frames can be acquired, thereby increasing N, reducing shot noise and improving the SNR of the detected PSF (**Fig. 3a**). The same effect is achieved by over-magnification of the image and subsequent spatial binning.

**Figure 3.**
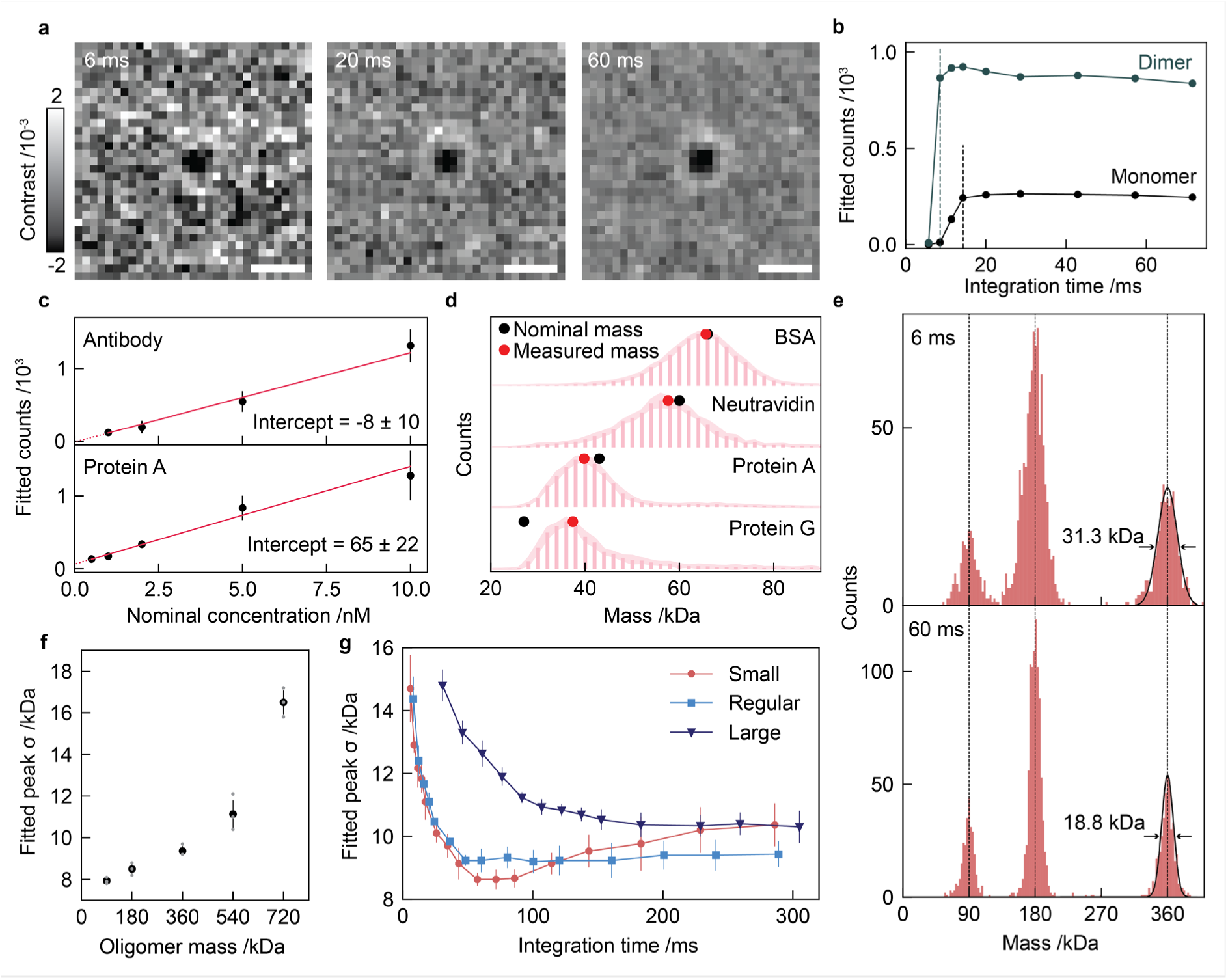
Shot noise and how it affects sigma and quantitative detection. (**a**) Effect of increased integration time on ratiometric image. (**b**) Detectability of Dyn1 ΔPRD monomer and dimer at arbitrary thresholds (**c**) Quantification of binding events for 150 kDa antibody (top) and 42 kDa protein A (bottom). (**d**) Nominal vs measured mass determination for BSA (66 kDa), neutravidin (60 kDa), Protein A (42 kDa) and protein G (27 kDa) calibrated to Dyn1. (**e**) Mass vs peak sigma for three repeats of Dyn1 ΔPRD large FOV. (**f**) Effect of increased integration time on Dyn1 ΔPRD histogram and tetramer FWHM. (**g**) Effect of integration time on Dyn1 ΔPRD tetramer sigma across standard FOVs.

Quantitative detection is critical for accurate characterisation of biomolecular affinities. An experimental approach to determine how quantitative detection is involves titrating towards zero analyte concentration: if detection is quantitative, a concentration vs counts plot should exhibit a linear dependence with an intercept near zero. For a high SNR protein such as a 150 kDa antibody at 100 ms averaging time, such behavior is indeed observed experimentally (**Fig. 3b**). Repeating the experiment for the same averaging time and analysis parameters for protein A (42 kDa), reveals a slight positive offset for the intercept, indicative of a small amount of noise features being counted for the given detection parameters (**Fig. 3c**). If required, one can iterate the acquisition and analysis parameters to optimise the performance as a function of analyte mass.

Signal strengths that approach the noise limit also influence mass accuracy in the low mass range. A low detection threshold will lead to substantial false positives, while a high detection threshold results in a loss of low SNR events. This will lead to an effective hard cut off on the low mass end of the distribution, thereby artificially shifting the average mass to higher values (**Fig. 3d**). This effect can be observed when characterising proteins in the 27 to 66 kDa mass range. As previously, this effect can be minimized by simultaneous optimization of averaging time and analysis thresholds, where required. In essence, this scenario will lead to an underestimate of the number of species and an overestimate of their mass, because only the high mass side of the distribution is detected. As a result, the mass-to-optical contrast relationship will drop off too slowly as lower mass species are detected.

Mass resolution between species of similar mass in a sample assay is influenced by many factors. Mass resolution is generally lower for larger species because of increased mass broadening due to local differences in glass roughness and the increasing difficulty to maintain measurement accuracy as the SNR increases substantially (**Fig. 3e**)^13^. This is a consequence of the difficulty with accurately estimating signals with SNRs>50 because even minor deviations in the fit used to extract the contrast can lead to errors. Mass resolution also depends on sufficient SNR to accurately estimate the individual peak contrast, with the full width at half maximum (FWHM) reducing from 31 kDa to 19 kDa with frame averaging for the tetramer peak of dynamin-ΔPRD shown here (**Fig. 3f**). The dependence of mass resolution on SNR can also be observed by its dependence on the size of the FOV, reducing *N* as optical power density is distributed over a larger area (**Fig. 3g**). Loss, rather than continuing gain of mass resolution for longer averaging times arises from additional, non-shot noise sources.

### Non-shot noise contributions

The results in **Fig. 3** demonstrate that several key performance parameters of MP can be improved by increasing the averaging time, which in turn reduces shot noise-induced background fluctuations. However, if this relationship were to hold indefinitely, there should be no limits to detection sensitivity or mass resolution for MP. We can test this relationship by computing signal fluctuations in the absence of analyte in both space and time. To do this, we evaluate the contrast standard deviation of a series of images on a pixel-by-pixel basis (**Fig. 4a**), which can be converted to a mass-equivalent using a mass-to-contrast calibration (**Fig. 1f**). For a purely shot noise-limited case, we would expect this noise to scale with the inverse square root of the averaging time.

**Figure 4.**
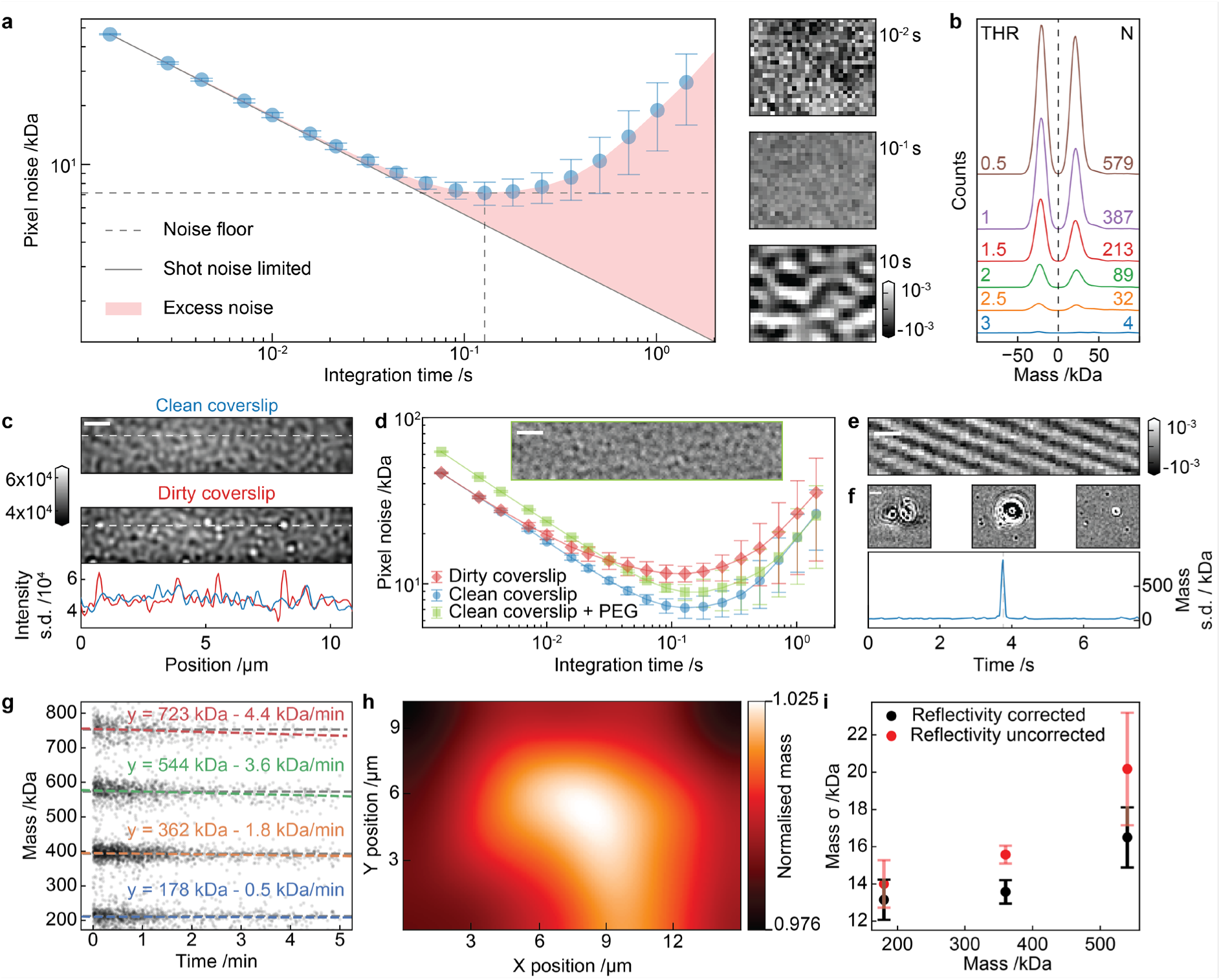
Noise contributions beyond shot noise. (**a**) Quantification of the median standard deviation over space and time for increasing integration times for a mass photometry video of PBS buffer. The difference between the datapoints (average ± standard deviation of 5 technical repeats) and the shot-noise limit as extrapolated from the first datapoint (solid line) constitutes excess noise (red). The noise floor (7.1 kDa) and corresponding integration time (128 ms) are indicated. Representative mass photometry images of the same field of view at three different integration times are shown. (**b**) MP histograms of a PBS video analyzed at 128 ms for various Welch’s T-test filter thresholds (‘Threshold 1’ in DiscoverMP, default = 1.5) for particle detection. (**c**) Representative native MP FOV for a PBS buffer on a clean and dirty coverslip. The plot shows the intensity standard deviation across the horizontal dashed line for both (clean coverslip in blue, dirty coverslip in red). (**d**) Same as (**c**), but now also for a dirty coverslip (red diamonds) and PBS buffer containing 0.5% (w/v) PEG 8K (green squares). Inset shows a representative field of view of a ratiometric mass photometry video of PBS buffer containing 0.5% (w/v) PEG 8K at an integration time of 128 ms. Contrast thresholds are as in (**a**). (**e**) Ratiometric mass photometry field of view of a buffer solution displaying fringes created by back reflections from the droplet. (**f**) Ratiometric mass photometry field of views of a large sample impurity binding the coverslip and continuing to wobble at the same position. Snapshots are separated by 200 ms. Timetrace shows the mass standard deviation over the field, with the landing event indicated by the vertical dashed grey line. (**g**) Mass of detected events over time for a mass calibrant protein measurement. A linear fit reveals a trend of a 0.5-4 kDa (or 0.5% of the mass) decrease per minute in the mean mass value. (**h**) Local mean map of contrast values for a single mass photometry peak across the field of view, showing a nonuniformity of around 5%, with strong spatial correlation. (**i**) Influence of the reflectivity correction option in DiscoverMP on the standard deviation of mass calibrant protein peaks. All scalebars are 1 μm.

With increasing integration time, the single image (**Fig. 4a**) pixel noise drops, following the expected 1/√N dependence, and reaches a minimum of 7.1 kDa at 128 ms (**Fig. 4a**, middle FOV). For longer integration times, a dynamic speckle-like background appears, and the pixel noise increases, deviating from the expected shot-noise limit **(Fig. 4a**, bottom FOV). The source of this excess noise is currently unknown, but we have previously ruled out sample drift and shown that excess noise cannot be removed by temporal averaging^12^. We have observed a similar noise limit across many different instruments, buffers, and integration times, suggesting that its source is connected to the measurement itself, rather than being the result of an imperfect experimental approach.

In practice, we have found that choosing an integration time about 20% shorter than the noise floor determined by this plot produces the overall best compromise between detection limit and mass resolution. Optimizing the integration time along with the correct threshold of minimum contrast change for a particle to be detected is critical to minimize false positive detection of low molecular weight species. Too high a threshold causes binding events of low mass species to go undetected, while a too low threshold leads to false positives.

In contrast to protein binding events, the spatiotemporal nature of the non-shot noise (**Fig. 4b**) results in its detection as an equal number of events on the positive and negative mass axis at the same mass. The number of events within these symmetrical peaks increases for a given integration time as the detection threshold is lowered (**Fig. 4b**). The appropriate threshold can be determined empirically by adjusting the analysis parameters to obtain <200 symmetric events in total when making a recording of pure measurement buffer prior to protein addition.

Having characterized the fundamental noise contributions associated with MP, we turn to the most common causes for deviations from optimal performance. A relatively trivial, but often encountered effect arises from sub-optimally cleaned microscope glass coverslips. Remaining impurities evidence themselves as high-contrast particles on the surface in the raw image (**Fig. 4c**). The signal generated by these large particles can easily saturate the camera leading to inaccurate mass quantification of any biomolecules landing on top of them. Pixel noise is also clearly elevated relative to a clean coverslip (**Fig. 4d**), likely due to the imperfect ratiometric removal of the large associated signals. Hence, we recommend moving to a different field of view without these large particles as evidenced by reduced image sharpness or repeating the coverslip cleaning procedure if a brief search for a clean area is unsuccessful. Increased background fluctuations can also be generated by macromolecules such as poly-ethylene glycol if present in high abundance in the buffer (**Fig. 4d**). The background fluctuations generated by these macromolecules results in correspondingly lower SNR (**Fig. 4d**) and increased false positive detection. In addition, back reflections from the water-air interface of the droplet can produce large fringes in the ratiometric field of view (**Fig. 4e**), which can be mitigated by adjusting solute volume and lateral position of the sample. Finally, depending on their purity, biological samples can contain large particles whose landing on the coverslip generates a signal well above the expected background signal (**Fig. 4f**). These can easily be excluded from the analysis as required by masking the affected regions out.

On top of these effects that can affect MP measurements generally, there are a few specific ones that contribute to a loss of mass resolution. An easily identifiable one is focus drift, which can occur when adding chilled analyte to the coverslip, which is at room temperature. A change in temperature leads to differences in local refractive index, affecting the optical contrast. Such behavior can be easily spotted and removed by plotting the contrast as a function of measurement time, mitigating unwanted mass broadening. In a standard landing assay, focus drift values are around 0.5% of the peak mass per minute (**Fig. 4g**). In some cases, non-uniform illumination and glass roughness can also cause local reflectivity variations which manifest themselves as differences in a local mass-to-contrast conversion. As a result, events in different parts of the field of view exhibit systematically different contrasts. This effect can be mitigated by only analysing parts of the field of view, comparing mass histograms from different sub-FoVs, or using reflectivity correction (**Fig. 4h, i**).

### Quality of surface binding

The fundamental concept of MP relies on accurately quantifying the change in reflectivity associated with a biomolecule binding to the glass-water interface. **Figs**. **3** and **4** illustrated the potentially detrimental influence of excessive background fluctuations caused by either shot noise or other sources. In addition, the dynamics of individual landing events themselves can cause an error in the extracted particle contrast.

The most commonly encountered protein dynamics near the interface observed by MP are illustrated in **Fig. 5a**. *Binding events* – the protein lands and remains stationary on the surface, indicating stable and optimal binding. *Unbinding events* – the protein temporarily adheres to the surface and then detaches. *Rolling events* – proteins that land on the surface followed by lateral movement, resulting in a distinct binary event with a binding head and unbinding tail in a ratiometric image. *Wobbling events* – proteins sometimes adsorb weakly to the surface and proceed to exhibit erratic, multidirectional movement. The latter two event types can sometimes be erroneously identified by the particle picking algorithm as multiple events.

**Figure 5.**
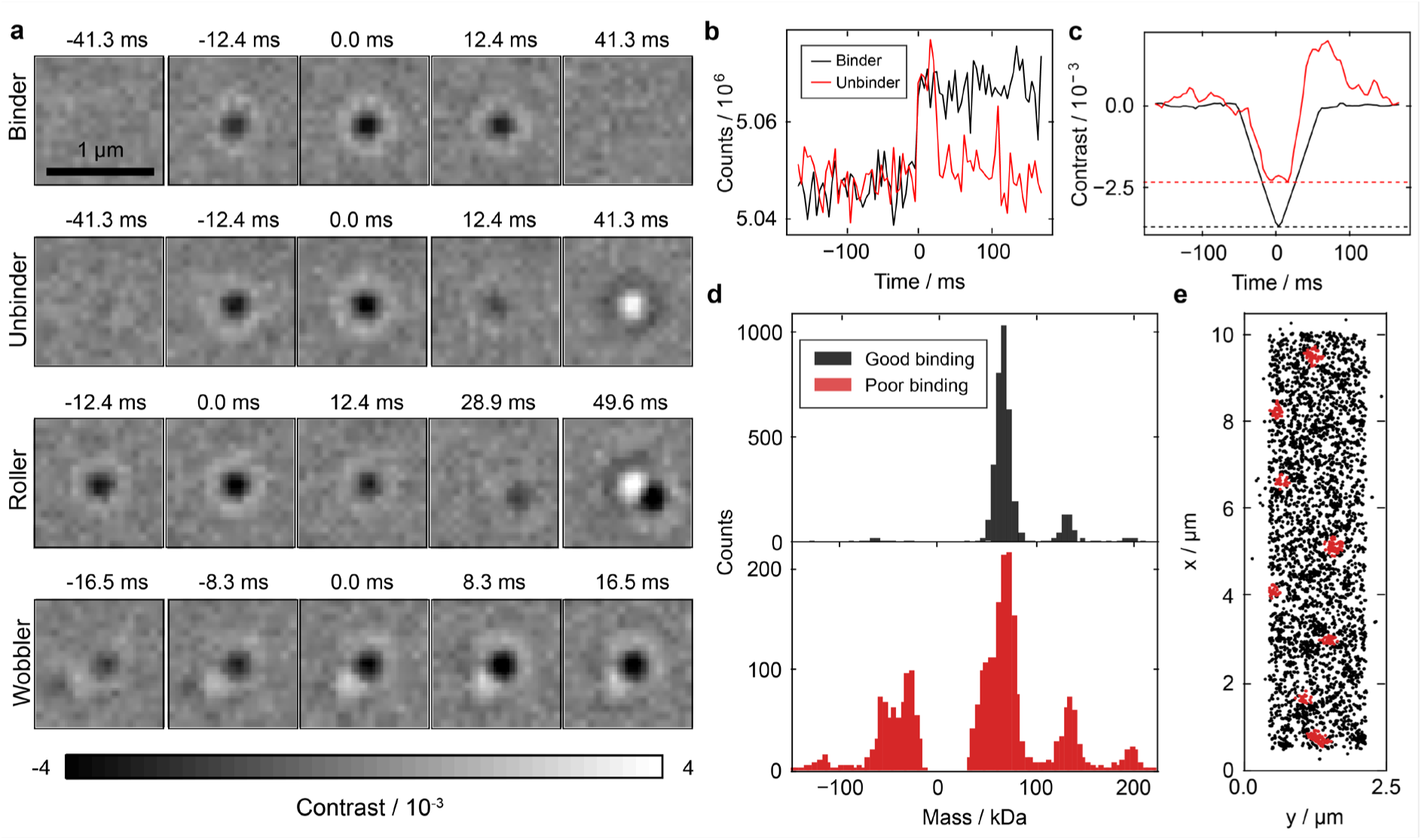
Accurate identification and quantification of protein events. (**a**) Series of frames depicting distinct types of landing events that may occur over the course a measurement. (**b**) Unprocessed time series traces showing a binding and an unbinding event for a BSA dimer. (**c**) Corresponding unbinding event ratiometrically processed over 80 ms integration. When compared with the average ratiometric trace of all optimal binders of the same species, a 34% reduction in contrast is observed for the singular unbinding event. This example illustrates the effect of poor binding on contrast estimation. (**d**) The accumulation of the effect of poor binding on a mass histogram of BSA. The good binding measurement used a new stock of BSA (black), the poor binding measurement used an old stock (red). (**e**) Nearest neighbour cluster filtering to remove overlapping/ wobbling events. Hotspots with abnormally high event density have been identified and highlighted in red.

Any of the characteristics that deviate from a regular binding event can be considered as a suboptimal event. This arises from the need to average frames to reduce shot noise: if the particle departs or moves during the integration time, it will decrease the local reflectivity change, causing a contrast and thus mass measurement error. The principle behind this effect is most easily understood when considering the consequences of rapid unbinding on the intensity of an individual camera pixel (**Fig. 5b**). Ratiometric processing of the single pixel trace of BSA dimers reveals a typical contrast of 0.35% when averaged over all optimally binding events (**Fig. 5c**). In the example with subsequent unbinding, however, the event yields a contrast of 0.23%, over a third less than expected. This contrast reduction is caused by the inclusion of the period without the particle present on the surface during the 80 ms integration time.

Although the impact of other suboptimal event types may be more subtle, one can easily understand how they lead to similar effects on the single (sub-diffraction) pixel level. It is their cumulative impact that can significantly distort the mass histogram (**Fig. 5d**) and manifests itself in several ways: The emergence of an unbinding peak, i.e., a symmetrically mirrored negative mass peak; the broadening of peaks, frequently accompanied by a characteristic shoulder on the low mass end; and an increase in baseline noise between peaks. These distortions complicate the interpretation of MP data, especially when coupled with additional noise sources, underlining the importance of accurately characterising and mitigating suboptimal binding events. A simple analysis-based approach to mitigate the effects of suboptimal binding is using a nearest neighbour-based clustering filter, which identifies hotspots of high event density, removing overlapping events and wobbling events (**Fig. 5e, red**).

An ideal measurement consists only of binding events. However, depending on the charge of the glass surface and the biomolecule of interest, binding can be reversible. In scenarios where the affinity of the biomolecule to the surface is low, binding, and unbinding events are observed even at low levels of surface coverage. This can lead to deterioration of measurement performance in terms of both mass accuracy and resolution (**Fig. 6a, b**). In general, the binding:unbinding ratio correlates well with key parameters such as linear scaling of total counts with analyte concentration, and the achievable mass resolution (**Fig. 6c**).

**Figure 6.**
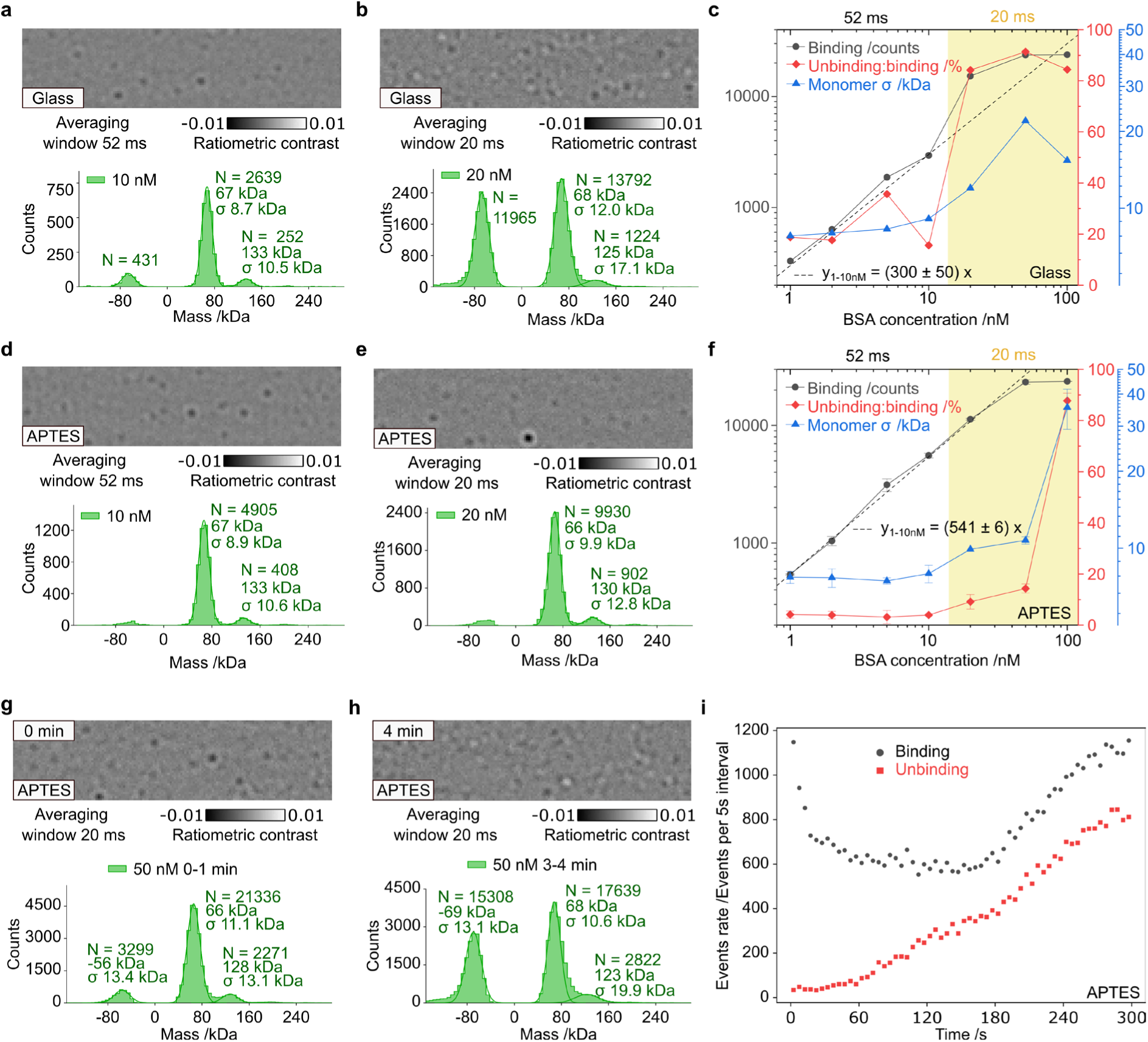
Effect of concentration and surface chemistry on precision, accuracy, and quantification. (**a, b**) Ratiometric frames and corresponding histograms from 1-minute measurements on glass at a concentration of 10 nM and 20 nM. (**c).** Dependence of binding counts, unbinding:binding ratio and monomer σ on BSA concentration on glass; slope was estimated from datapoints in concentration range of 1-10 nM. (**d, e)** Frames and histograms of BSA on APTES at a concentration of 10 nM and 20 nM. (**f)** Binding counts, peak ratios and monomer σ as function of BSA concentration on APTES, slope was estimated in concentration range of 1-10 nM. (**g, h)** First and fourth minute of measurement at 50 nM concentration and (**i)** events rates evolution during 5 mins.

For BSA, functionalising the microscope cover glass surface with positively charged (3-Aminopropyl)triethoxysilane (APTES) increases the surface affinity, evidenced by the doubling of molecular counts for the same analyte concentration and a drop in unbinding counts (**Fig. 6d**). The increase in protein binding to the surface improves mass resolution at the same analyte concentration but also enables higher measurement concentrations because of reduced unbinding events that would otherwise contribute to overcrowding of field of the view (**Fig. 6b, e**). The corresponding titration exhibits similar resolution and slightly improved count scaling with concentration in the 1-10 nM analyte concentration range. In the 10-50 nM concentration range, however, APTES functionalization markedly improves those parameters over those achievable on a bare glass surface. APTES functionalization thus improves the upper concentration limit in this case about five-fold with overall identical performance (**Fig. 6f**). At higher concentrations, e.g., 100 nM, the measurement fails due to overcrowding of the field of view and saturation of the surface, even on APTES.

The effect of protein surface saturation can also manifest itself during the measurement, as more proteins bind to the surface. For example, at 50 nM, low levels of unbinding are observed during the first minute of measurement (**Fig. 6g**). After 3 minutes, however, the APTES surface becomes saturated, and a substantial unbinding peak emerges (**Fig. 6h**). The temporal evolution of binding and unbinding demonstrates that optimal binding conditions are satisfied for only the first minute of the measurement, beyond which the measurement quality degrades (**Fig. 6i**). These results illustrate the advantage of starting the measurement immediately after sample addition to avoid unnecessary surface coverage by the analyte.

### Accurate quantification of oligomeric ratios including <100 kDa species

Whether an analyte can be detected by MP depends on both the averaging time and its mass, because the two of them define the signal-to-noise-ratio. As event contrast scales linearly with species mass, it is intuitive that the SNR decreases for smaller species, which can be compensated with longer averaging time, of particular importance when one of the species is in the <100 kDa range (**Fig. 3b**).

Increasing the averaging time leads to improved detection at low mass, which directly translates into a more accurate monomer:dimer ratio as shown here for BSA (**Fig. 7b**). Increasing the averaging time further may improve mass resolution but has no impact on the total number of detected events or the monomer:dimer ratio. This suggests that the peak ratio is quantitative for averaging times >20 ms.

**Figure 7.**
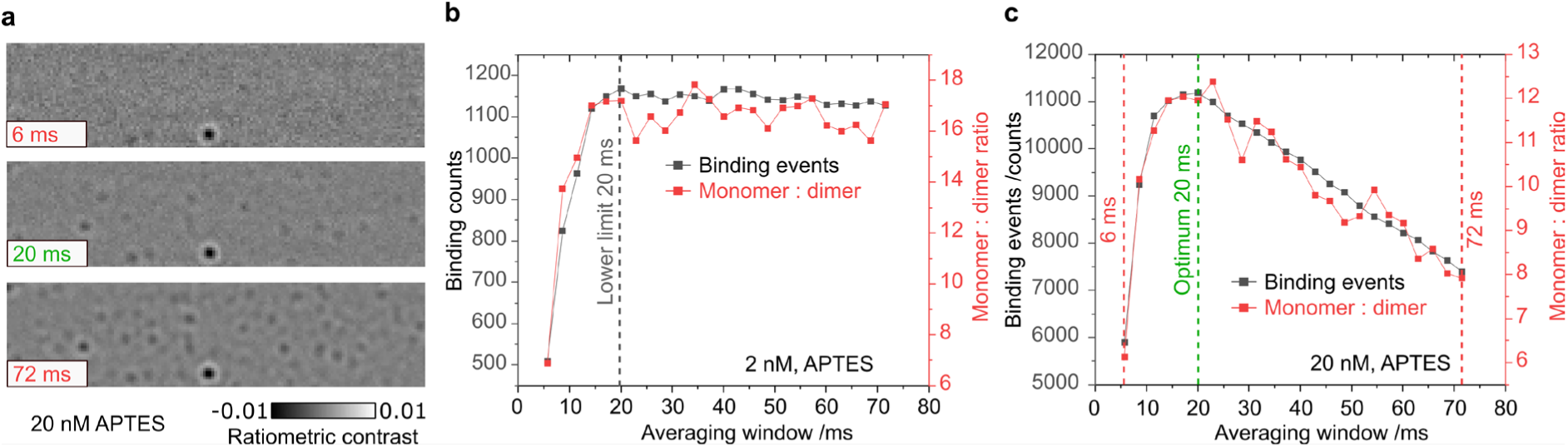
Effect of concentration and averaging window on quantifications of oligomer ratios. (**a**) Same ratiometric frame at concentration of 20 nM processed using different averaging windows of 6, 20 and 72 ms. (**b**, **c**) Number of detected binding events and monomer:dimer peaks ratio in dependence on averaging window at a concentration of 2 nM and 20 nM BSA.

At 10-fold higher analyte concentration, however, longer averaging times lead to a drop in total detected events and the monomer:dimer ratio (**Fig. 7c**). Event crowding makes it more challenging to detect low contrast events in the presence of larger species, an effect that becomes more prominent with increasing averaging time. This, together with the need for sufficient averaging time to detect weak events leads to clear maximum of both the monomer:dimer ratio and number of detected events at the optimum averaging window of 20 ms. The fact that this measurement is indeed quantitative is indicated by the observation that the total detected counts are 10-fold higher than those observed at 2 nM. In addition, the monomer:dimer ratio drops from 17 to 12 as expected for a higher measurement concentration. We emphasize that total counts can indeed be quantitative and scale with analyte concentration, with the slope error only of 1.1%, when taking care with sample addition and using a good binding surface (**Fig. 6f**).

### Importance of concentration on measurement quality

The previous sections have shown the influence of protein concentration and the resultant landing density that appears in ratiometric processed images. To summarise, too high concentration can cause issues with:

- Protein binding: Saturated surfaces reduce binding quality, resulting in unbinding, wobbling, and rolling molecules.
- Quantitative detection: Overlapping PSFs will not be reliably detected, reducing molecular counting accuracy.
- Contrast accuracy: Overlapping signals cause errors in contrast estimation.
- Mass sensitivity: Overlapping PSFs are exacerbated at high averaging times. Increasing averaging to detect lower mass species will be ineffective.

As the effective landing density is also a function of protein oligomerisation, one cannot define a single concentration that can be used for every protein. Instead, what matters is the event density, which can be adjusted for optimal performance by titration for optimal mass resolution and mass accuracy.

### Overview of Procedure

To address potential shortcomings that can be encountered when making an MP measurement we provide a protocol that addresses the key areas that affect measurement quality; surface quality, mass calibration, mass measurement and, data analysis (**Fig. 8**). To demonstrate the efficacy of the protocol we detail experimental conditions for two proteins, BSA and SARS-CoV-2 antibodies. BSA as an example of a poor binding low mass oligomeric protein, which is challenging to measure with high accuracy and reproducibility and the SARS-CoV-2 antibody due to the community interest in using MP as a technique to characterise antibody-antigen affinities. The provided procedure and concentrations are to be used as a starting point for measuring your protein of interest, the same optimization strategy can be applied to any sample and is intended to push the limits in terms of sensitivity, resolution, and quantification.

**Figure 8.**
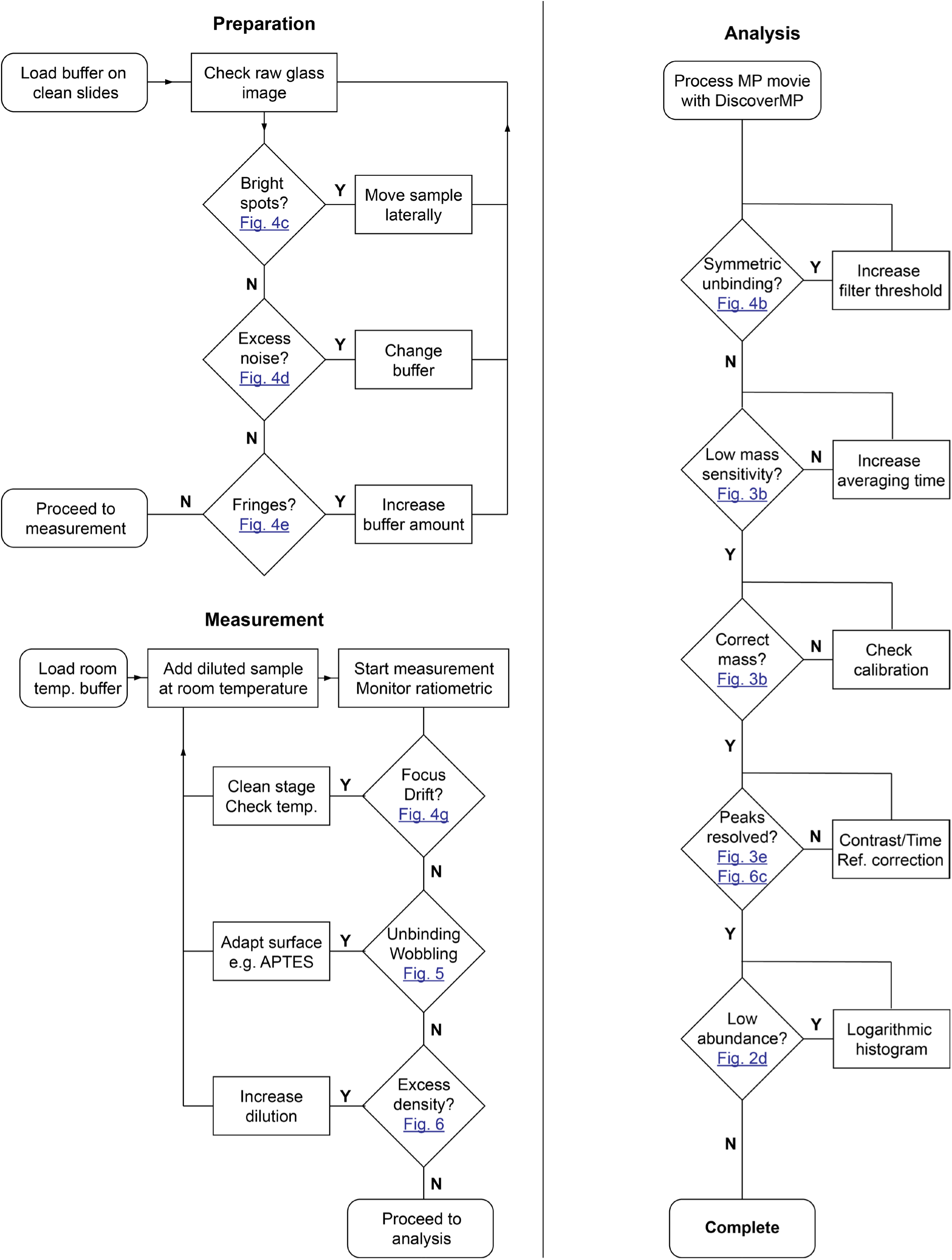
Flow diagram for how to do the best possible MP measurement. The four areas covered are slide preparation, mass calibration, sample measurement and analysis. Example troubleshooting suggestions are given for each step where relevant and is intended as an exhaustive description of all potential issues that could be present in an MP measurement. The typical measurement will not experience all these issues, but the user should be mindful of these potential noise sources which can reduce measurement quality.

### Expertise needed

To perform MP, basic knowledge of how to use a mass photometer using the AcquireMP software and how to perform analysis using the DiscoverMP software is required. Anyone with experience with standard wet lab techniques will be able to perform the remainder of the protocol without difficulty.

#### Limitations

1. **Detergents**: Detergent micelles in themselves are protein-sized objects and thus produce comparable signal that creates substantial background. This can be mitigated using special detergents, or peptidiscs^14,15^.
2. **Protein mass:** The protein mass and the mass differences between biomolecules complexes need to be larger than 30 kDa for quantitative measurements.
3. **Buffer and protein purity**: Proteins and buffer will ideally be free of any other particles. High concentrations of low mass particles, such as PEG or glycerol below the detection limit can cause additional noise. Note that buffer ions are sufficiently small to not contribute to background noise.
4. **Low-affinity interactions**: Complexes or oligomers can break down at low concentrations required by MP. Therefore, complexes with an off-rate on the order of seconds and a dissociation constant above 100 nM cannot be detected in a standard landing assay. This can be overcome by measurements using surfaces allowing measurements at high concentrations in equilibrium or exploiting rapid dilution using microfluidics^16,17^.

### Materials

#### Reagents

- Gibco™ DPBS, no calcium, no magnesium (Fisher Scientific, cat.no. 15326239)
- 3-Aminopropyltriethoxysilane 99% corc-sealed (Sigma-Aldrich, cat.no. 440140)
- acetone 99.8%, HPLC grade (Sigma-Aldrich, cat.no. 270725)
- 2-propanol ACS reagent, ≥99.5% (Sigma-Aldrich, cat.no. 190764)
- Bovine serum album lyophilized powder (Sigma-Aldrich, cat.no. B6917)
- SARS-CoV-2 (2019-nCoV) Spike Neutralizing Antibody, Mouse Mab (SinoBiological, cat.no 40591-MM48)
- Massference-p1 (Refeyn MP-CON-41033)
- MilliQ water, >18 MΩcm, 0.22-μm filtered, ultrapure (type 1) water
- Nitrogen N5.5 99.9995% Research Grad (Boc)

#### Equipment

- TwoMP (Refeyn)
- DiscoverMP Version 2023 R1.2 (Refeyn)
- *For using Refeyn consumables*
- Sample Preparation Kit (*Optional*, MP-CON-21008, Refeyn)
- Samples well cassettes (6 wells) for the OneMP, TwoMP, SamuxMP (MP-CON-41005, Refeyn)
- *For cleaning own coverslips*
- Carbon PEEK Replaceable Tip Tweezers (Agar Scientific, cat.no. AGT551)
- Fisherbrand™ Borosilicate Glass Tall Form Beakers 400ml (Fisher Scientific, cat.no. 15439093)
- Epredia No 1.5 coverslips 50 x24mm (VWR, cat.no. 16002-264)
- Slidebox for 25 Slides (GLW, cat.no. K25W)
- Optical Grade Cotton-Tipped Applicators (Thorlabs, cat.no. CTA-10)
- Carl Zeiss Immersol 518 F Immersion Oil (Fisher Scientific, cat.no. 10539438)
- Zepto Plasma Cleaner (Diener Zepto-BRS 200)
- Epredia™ Coverglass Staining Rack (Fisher Scientific, cat.no. 12627706)
- Ultrasonic Cleaning PROCLEAN 10.0S (Ulsonix)
- Lens Cleaning Tissue (Thorlabs, cat.no. MC-5)
- Velp Scientifica™ AREC.X 7 Digital Ceramic Hot Plate Stirrer (Fisher Scientific, cat.no. 17274083)
- Magnetic stirrer bar, 25 mmx 8mm (Sigma, cat.no. HS120549)
- RS PRO Infrared Thermometer (RS, cat.no. 136-7890)
- B Braun™ Injekt Solo Cone Syringes (Fisher Scientific, cat.no. 12722637)
- B Braun™ Sterican 70mm 20-gauge needle (VWR, cat.no. 612-0159)
- B Braun™ Sterican 16mm 25-gauge needle (VWR, cat.no. 612-0153)
- Silica Gel (RS, cat.no. 388-8421)
- Eppendorf® PCR tubes (Sigma-Aldrich, cat.no. EP0030124537)
- Eppendorf® Flex-Tubes Microcentrifuge Tubes (Sigma-Aldrich, cat.no. EP02236411)
- SalvisLab Vacucenter Vacuum Oven (Cole Parmer, cat.no. WZ-52402-36)

### Reagent setup

#### Massference-p1 preparation Timing 30 minutes

1. Defrost a vial of Massference-p1
2. Aliquot into 2 µl vials.
3. Freeze at -20°C and store until required.

#### BSA preparation Timing 30 minutes

1. Prepare a 60 µM solution in Dulbeccos PBS.
2. Store at 4°C until measurement, can be stored for up to 1 month.

### Antibody preparation

#### Timing 3 minutes

1. Prepare an 8.3 µM solution in Dulbeccos PBS.
2. Store at 4°C until measurement, can be stored for up to 1 month.

### Monthly instrument calibration: Timing 2.5 hours

- Before a large set of experiments, or in one-month intervals, follow the instrument manual to calibrate the MP device acquisition image.

1. Switch on the mass photometer 2 hours prior to measurement.
2. Place an array of six gaskets onto the center of cleaned coverslip.
3. Put a drop of oil onto the objective and place coverslip onto the sample holder. Ensure sufficient and not excess oil.
4. Add 20 µL of MilliQ in the gasket, wait for 1 minute with a closed lid to equilibrate temperature.
5. Focus on the coverslip
6. Got to ‘Acquisition’ in the menu bar in the top of the screen and select ‘calibrate the acquisition image’.

### Procedure

#### Coverslip Cleaning (If using precleaned slides from Refeyn, go to protein measurement): Timing 30 minutes

1. Place coverslips in a cleaning rack with tweezers or gloved hands, avoid touching the measurement area.
2. Clean coverslips by sonicating in acetone, 1:1 MilliQ:Isopropanol solution and MilliQ, each for 5 minutes.
3. Blow dry coverslips with nitrogen and store them in a box. **TROUBLESHOOTING**

#### Surface Amination (if needed): Timing 1.5 hours

1. Pour 250 mL of HPLC acetone into a beaker, cover it with aluminum foil to avoid solvent evaporation and heat the solution to 50°C (measured by an optical thermometer). **Critical Step.** Low-water content acetone is important to avoid APTES polymerization and aggregate formation, making measurements of small proteins impossible.
2. Add 5 mL of APTES to the beaker, using a syringe with an attached needle, to obtain a 2% solution.
3. To activate the glass surface with -OH groups for the following silanization, insert cleaned coverslips into the oxygen plasma cleaner.
4. Pump the chamber to a pressure below 0.15 mbar.
5. Introduce O_2_ to reach 0.8 mbar pressure.
6. Plasma-clean the coverslips for 8 minutes at 50% power.
7. After plasma cleaning, promptly put the coverslips into a beaker with acetone (99.8% HPLC grade), and then immediately transfer into the beaker with 2% APTES solution.
8. Incubate coverslips with APTES at 45°C for 15 min on magnetic stirrer.
9. Sonicate APTES beaker with coverslips for 1 minute to remove physiosorbed APTES molecules and aggregates.
10. Incubate for another 15 minutes while stirring at 45°C. Sonicate the slides in fresh acetone twice for 5 minutes, replacing the acetone between each sonication and then sonicate for 1 minute in MilliQ **TROUBLESHOOTING**.
11. Dry APTES coated slides with nitrogen and store in a box with silica gel.
12. Clean APTES beakers by removing the 2% APTES solution and sonicating in fresh acetone for 30 minutes.

- **Critical step.** Quickly transferring the coverslips from the plasma cleaner to the APTES beaker is essential as -OH group activation decreases with time.
- **Pause Point:** For best performance, use the slides the day of preparation but they can be stored and used for 1 week; For some proteins, such as BSA, the performance does not substantially depend on storage duration.

### Protein measurement

#### Measurement Preparation: Timing 2 hours

1. Ensure microscope objective is completely clean before starting. Clean with IPA and optical tissues and cotton buds if not.
2. Switch on the mass photometer 2 hours before measurement to ensure thermal equilibrium.
3. Prepare an Eppendorf with a 1.5 mL aliquot of buffer and equilibrate buffer at room temperature.
4. If using home-made slides, place six gaskets onto the center of a coverslip with the same surface chemistry as for the measurement, e.g., APTES functionalized glass.
5. Set the machine to advanced mode and frame binning 2.
6. Select FOV according to intended analyte.

a. Small for <70 kDa (e.g., BSA).
b. Regular for >70 kDa (e.g., SARS-CoV-2 antibody).

#### Buffer background measurement: Timing 5 min (for the first measurement in a set of gaskets)

1. Fill the gasket with 10 µL buffer and focus using AUTOFOCUS. You can check that optimum focus has been found by manually moving the Z stage to verify that the image sharpness is at a global maximum. **TROUBLESHOOTING**
2. Check the native image for bright spots which saturate the camera, reposition the slide laterally if these are present. Sharpness should be less than 6%.
3. Check the ratiometric image for any landing events or fringes, see troubleshooting if you observe either of these.
4. Record a movie for 60s. **TROUBLESHOOTING**
5. Optional: By moving the Z stage verify that optimum focus held during the measurement if not, repeat steps 7-12.

#### Calibration: Timing 10 minutes

6. Immediately after buffer measurement, add to the same well 10 µl of 250x diluted Massference-p1 for a total 500x dilution, aspirating 3 times with the pipette.
7. Check the focus position is still the same, if not refocus and record a 60s movie.
8. Process the movie with standard analysis settings in DiscoverMP. Fit gaussians to the 86, 172, 258 and 344 kDa peaks.
9. Process movie of buffer only using same analysis in DiscoverMP and apply calibration. Check the histogram for buffer movie, there will typically be <30 binding events total in a clean buffer movie for averaging time of 20 ms and default threshold. TROUBLESHOOTING.

- **Optional step**: Repeat buffer measurement and calibration every hour, or every second slide, or after last measurement. If buffer movie background is increased over previous measurement, prepare another 1.5 mL Eppendorf with buffer.

#### Sample measurement: Timing 5 minutes/measurement

1. Dilute stock solution to working solutions.

a. 60 µM BSA stock stored at 4 to 12.5 µM. Keep 12.5 µM solution on ice during the measurement.
b. 8.3 µM SARS-CoV-2 antibody to 6.25 µM, keep on ice during the measurement.
2. Dilute 12.5 µM BSA to 12.5 nM of volume 20 µL and incubate at room temperature for 2 minutes.
3. During incubation period, change to a fresh gasket on a slide, add 4 µl of buffer and find focus. **TROUBLESHOOTING**
4. Find an area of the slide with no saturation spots in the native image, check ratiometric movie for landing events or fringes and refocus on the spot intended for measurement.
5. Add 16 µl of protein to the 4 µl of buffer in the gasket, for a total concentration of 10 nM, aspirate 5 times to mix the solution.
6. Close the lid and start acquisition immediately, recording for 60 s.

- **Critical step.** For reproducible measurements, the incubation time in the Eppendorf should be as similar as possible between repeats. Ensure that the sample is at room temperature before adding to the gasket. Add buffer 2 minutes before measurement to avoid evaporation. For working at concentrations <1 µM use PCR tubes instead of standard Eppendorf tubes to reduce protein adsorption
- **Critical step**: For best performance, acquire 3 repeats at 10 nM and perform titration at final concentrations of 2, 5, 10, 20, 50 nM in the gasket. In the case of analytes with long dissociation rates incubate at 2, 5, 10, 20, and 50 nM concentration.

### Analysis: Timing 10 minutes per measurement

1. Process BSA/Antibody movies using the same analysis settings as used for calibration and buffer background estimation.
2. Apply the mass calibration from the Massference-p1 measurement.
3. If the 66 kDa peak of BSA is not observed, increase the integration time by changing the number of averaged frames in the measurement settings until it fully appears.
4. Check the negative mass side of the histogram, if there is a symmetric unbinding peak at low mass, increase Threshold 1 until the total counts in the positive and negative low mass range are below 200.
5. Fit the monomer and dimer peaks, the mass should be 66 kDa and 132 kDa with a sigma <10 kDa for the BSA monomer. Expected mass for SARS-CoV-2 antibody should be 150 kDa with sigma of < 10 kDa. **TROUBLESHOOTING**
6. Compare the monomer counts between all the peaks for each technical repeat, fluctuations in counts should be within 25%, ideally <15 %, when added carefully and in a reproducible fashion.
7. If measured analyse the measurements at 2, 5, 10, 20 and 50 nM using the same analysis parameters.
8. Plot the binding counts. They should increase linearly with concentration (**Fig. 6 c, f**, and **Fig. 9c**), slope error in the optimum concentration range should be < 5%, ideally < 2%. **TROUBLESHOOTING**
9. Plot monomer peak sigma and binding:unbinding ratio as a function of concentration. Both parameters should be constant below 20 nM (**Fig. 6 c, f**). **TROUBLESHOOTING**
10. Export the fitted events from DiscoverMP and plot the contrast/mass against the frames indices, contrast/mass should be stable over time (**Fig. 1f**).
11. To determine if a specific concentration and chosen integration time is suitable for estimation of accurate oligomer ratios, repeat the analysis with different integration times by pressing SETTINGS and using increasing the averaged frames to from 3-20 in single frame steps. Plot the BSA monomer/dimer ratio with respect to integration time (**Fig 7 b, c**).

### Trouble Shooting

See **Table 1** for troubleshooting.

**Figure 9.**
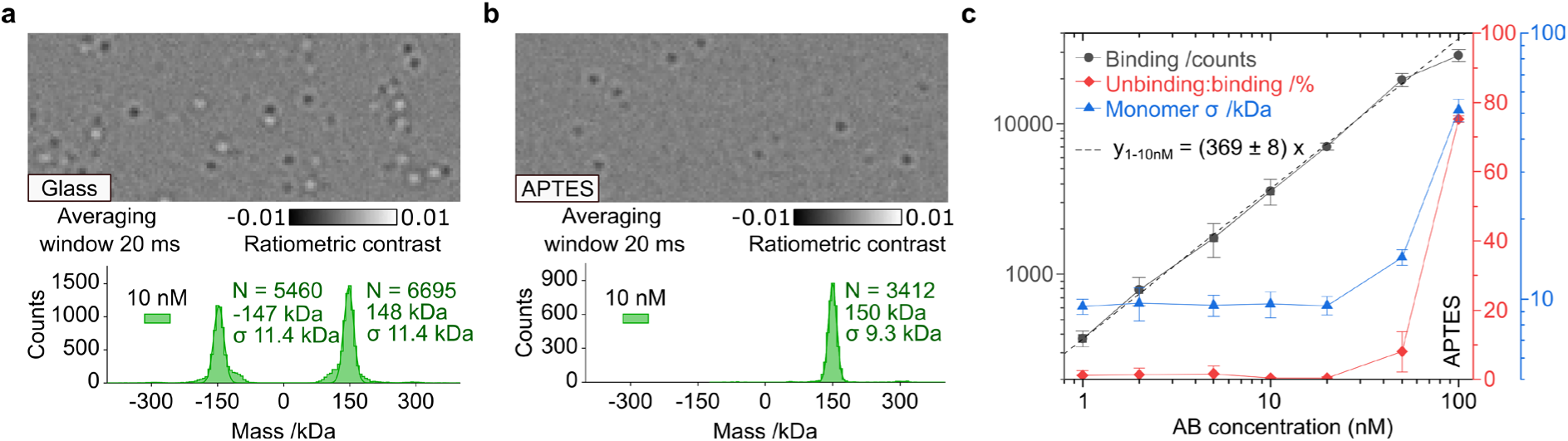
Titration measurement of SARS-CoV-2 antibodies. (**a)** Measurement at 10 nM on glass. **(b)** Measurement at 10 nM on APTES. **(c)** Concentration dependence of counts, unbinding/binding ratio and monomer σ, which was measured on APTES surface; the slope was estimated in the concentration range of 1-10 nM.

**Table 1.** Troubleshooting. Additional troubleshooting suggestions can be found in the user manuals of AcquireMP & DiscoverMP.

| Step | Problem | Reason | Solution |
| --- | --- | --- | --- |
| Coverslip cleaning 3 | Dirty glass after cleaning | Dirty glassware | Clean glassware with glass detergent such as Hellmanex, rinse well with DI water |
|  |  | We sometimes observe that for a pack of coverslips one side is always consistently cleaner than the other. The source of this variation is unknown | Use the other side of the coverslip |
| Surface Amination 10 | Aggregates on APTES slides | APTES has been overly exposed to moisture from the air | Replace with fresh APTES |
|  |  | Contamination of glassware with APTES aggregates | Always keep one specific beaker for APTES functionalisation, sonicate for 30 mins in acetone after functionalising. |
|  |  | APTES stock solution contains aggregates, that developed over time. | Use new APTES stock bottle. Alternatively, reduce incubation time with APTES to 1 min, do not sonicate after silanization, just twice wash by agitation for 1 min in beaker with fresh Acetone. Then nitrogen blow dry coverslips and crosslink APTES in the oven at 110°C set for 2 hours. Then perform standard coverslip cleaning procedure. |
| Buffer Background 1 | Gaskets leaking on liquid addition | Gaskets have become dirty | Replace set from the pack, always keep the gaskets covered with the film. |
| Buffer background 4 | High numbers of events in buffer | Buffer is contaminated | Replace with new buffer, filter sterilise with 0.2 um filter. |
|  |  | Buffer has formed aggregates | Reduce concentration of any PEG, detergent etc. |
| Calibration 4 | Focus is not stable during measurement | Focus ring not sufficiently continuous due to bubbles | Lift the coverslip on one side and allow bubbles to move to the side then |
|  |  | Oil on the stage | Remove the coverslip, clean the stage with cotton buds and IPA. Change to new coverslip. |
|  |  | Protein solution is not at room temperature | Wait longer for the solution to equilibrate to room temperature then add to the coverslip |
|  |  | Protein buffer not identical to buffer loaded for focusing | Always use the exact same protein buffer to find focus as for the measurement. |
|  |  | There was low volume of liquid in the gasket used for focusing | Find the minimum volume for focusing by incremental addition of liquid by 1 µL, minimal volume when focus and image properties stay the same the after addition. |
| Sample measurement 4 | No events detected. | Protein was deposited on the tube walls or pipette tips during dilutions and mixing. | Use PCR tubes, mix quickly by aspirating up and down, reduce incubation times at low concentrations. |
|  |  | Protein is aggregated. | Avoid making bubbles during dilutions avoid fast vortexing. |
| Analysis 5 | Mass is inaccurate. | Inaccurate mass calibration | Check that the correct contrast has been assigned to each peak. Check that the mass to contrast slope is linear. Repeat measurement until $R^2 > 0.96$ is achieved. |
|  |  | Touching the coverslip during adding protein and changing the focus position | Take care not to touch the coverslip, you can pipette from a reasonable distance away from the gasket. The autofocus can only correct for a limited shift in Z, check the glass is in focus before recording and refocus if necessary |
| Analysis 7 | Contrast/mass is inconsistent between measurements. | Incorrect focus position being used. | If using the auto find focus features, check that the highest sharpness is found by moving the stage manually up and down. |
|  |  | Insufficient warmup time for the instrument to equilibrate. | Leave the instrument on longer prior to the measurement. Take regular calibration measurements during the day to account for other contrast drift sources. |
|  |  | Different buffer used for measurement, or method of protein addition changed the focus position. | Manually fine refocus Z stage right before acquiring each movie. |
| Analysis 8 | Counts are inconsistent between measurements. | Measuring repeatedly from protein solution at low concentration. | Make a fresh dilution prior to each measurement to avoid proteins sticking to the Eppendorf side. |
|  |  | Inconsistent time between adding protein and pressing record. Most of the protein will immediately bind to the glass and the binding rate decays rapidly as proteins are depleted from solution | Always try to leave the same amount of time between adding the protein and recording the measurement |
|  |  | Insufficient mixing when adding protein solution to droplet in the gasket. | When adding the protein, mix well by aspirating several times in the gasket. |
| Analysis 9 | Mass peaks always broader than stated values | Incorrect focusing position | Check sharpness is maximized for each measurement at selected spot for measurement. |
|  |  | Particle concentration too dense | Ensure dilutions are correct |
|  |  | Poor binding to surface | Check in the negative part of the histogram for number of unbinding molecules, functionalise new set of APTES slides with fresh APTES |

### Anticipated Results

Massference-p1 at 500x dilution from stock concentration exhibits peaks at 86, 172, 258 and 344 kDa (**Fig. 1g**). Additional peaks can be observed beyond the 344 kDa peak but there is limited additional benefit to the calibration by including them. The expected sigmas for these fitted peaks are 10, 13, 17 and 17 kDa respectively, we have observed that baseline resolution is not achieved at this concentration, but for the purposes of mass calibration the peak resolution is adequate. The calibration curve shows a mass to contrast ratio of -0.03 1/kDa, which is typical for a well performing instrument, the R^2^ of the mass contrast fit is 0.9997, measurements below 0.95 should be repeated an R^2^ above this is achieved. Baseline resolution of BSA monomer and dimer is achieved at 10 nM on an APTES surface, with mass measurements of 66 and 132 kDa for the monomer and dimer respectively (**Fig. 6a**). Measurement at this high limit concentration is important for precise affinity estimates of weaker protein-protein interactions^5^. SARS-CoV-2-antibodies at 10 nM show high unbinding on glass and broadening of the peak due to inaccurate contrast estimation of poor binding events (**Fig. 9a**). Unbinding is effectively eliminated on APTES, allowing access to higher concentration ranges and consequently lower affinity interactions (**Fig. 9b**). The measured counts are linear with increasing concentration up to 50 nM and sigma <10 kDa can be obtained at all these concentrations, giving a resolving power >16 for measuring antibody antigen interactions (**Fig 9c**).

## Author Contribution Statement

J.K, R.W, D.L, JC.T, J.B, K.I are co-first authors, they contributed equally, everyone agrees their authorship orders can be exchanged to benefit their own career development. S.T and P.K are joint corresponding authors. All authors contributed to the conceptualisation. Data acquisition, analysis, interpretation, and writing of corresponding protocol section: Fig 1 JC.T; Fig 2, 3d, 4g-i D.L; Fig 3a-c,e-g J.B; Fig 4a-f R.W; Fig 5 K.I; Fig 6, 7, 9 JK; Fig 8 S.T. Writing of introduction S.T, review and editing of final manuscript S.T. and P.K. Supervision S.T and P.K.

## Acknowledgements

This work was funded by the European Research Council (ERC) Consolidator Grant PHOTOMASS 819593(P.K, K.I), the Engineering and Physical Research Council (EPSRC) Leadership Fellowship EP/T03419X/1 (P.K, JC.T), the Biotechnology and Biological Sciences Research Council BB/W00349X/1, (P.K, S.T), the Wellcome Trust, Grant Number: 218514/Z/19/Z (R.W), UK Research and Innovation (UKRI) under the UK government’s Horizon Europe funding guarantee through project Marie Skłodowska-Curie Actions (MSCA) Postdoctoral Fellowship NanoMassCreator (101062868) EP/X025713/1 (J.K), Clarendon scholarship, Menasseh Ben Israel scholarship Kingsgate scholarship (D.L), EPRSC Doctoral Training Partnership (J.B).

For the purpose of Open Access, the author has applied a CC BY public copyright licence to any Author Accepted Manuscript (AAM) version arising from this submission.

The authors thank Konstantin Zouboulis, Francesca Naughton-Allen, Alexander O’Shea for feedback on the manuscript and discussion. The authors thank Manish Kushwah for providing the Dyn-1 protein. The authors also thank Refeyn for providing the Massference-P1 calibration protein and for providing useful feedback on the manuscript.

